# Shared and unique consequences of Joubert Syndrome gene dysfunction on the zebrafish central nervous system

**DOI:** 10.1101/2024.02.15.580456

**Authors:** Alexandra R. Noble, Markus Masek, Claudia Hofmann, Arianna Cuoco, Tamara D. S. Rusterholz, Hayriye Özkoc, Nadja R. Greter, Nikita Vladimirov, Sepp Kollmorgen, Esther Stoeckli, Ruxandra Bachmann-Gagescu

**Author notes:** **Correspondence:** Ruxandra Bachmann-Gagescu, Department of Molecular Life Sciences, University of Zurich, Winterthurerstrasse 190, 8057 Zurich. equal contribution (Alexandra R. Noble, Markus Masek).

## Abstract

Joubert Syndrome (JBTS) is a neurodevelopmental ciliopathy defined by a highly specific midbrain-hindbrain malformation, variably associated with additional neurological features. JBTS displays prominent genetic heterogeneity with >40 causative genes that encode proteins localising to the primary cilium, a sensory organelle that is essential for transduction of signalling pathways during neurodevelopment, among other vital functions. JBTS proteins localise to distinct ciliary subcompartments, suggesting diverse functions in cilium biology. Currently, there is no unifying pathomechanism to explain how dysfunction of such diverse primary cilia-related proteins results in such a highly specific brain abnormality. In order to identify the shared consequence of JBTS gene dysfunction, we carried out transcriptomic analysis using zebrafish mutants for the JBTS-causative genes *cc2d2a^uw38^, cep290^fh297^, inpp5e^zh506^, talpid3^i264^* and *togaram1^zh510^*and the Bardet-Biedl syndrome-causative gene *bbs1^k742^.* We identified no commonly dysregulated signalling pathways in these mutants and yet all mutants displayed an enrichment of altered gene sets related to central nervous system function. We found that JBTS mutants have altered primary cilia throughout the brain, however do not display abnormal brain morphology. Nonetheless, behavioural analyses revealed reduced locomotion and loss of postural control which, together with the transcriptomic results, hint at underlying abnormalities in neuronal activity and/or neuronal circuit function. These zebrafish models therefore offer the unique opportunity to study the role of primary cilia in neuronal function beyond early patterning, proliferation and differentiation.

**Summary Statement:** Joubert Syndrome gene dysfunction in zebrafish leads to abnormal brain cilia, altered transcription of neuron-associated genes and abnormal swimming behaviour despite normal brain morphology.

## Introduction

Ciliopathies are a diverse group of disorders arising from primary and motile cilia dysfunction. Primary cilia are small, sensory organelles present on almost all vertebrate cells. Ciliary dysfunction can therefore affect multiple tissues and organs (Reiter and Leroux 2017). In particular, many ciliopathies present with neurological abnormalities, underlining the crucial role of primary cilia during development and function of the central nervous system (CNS) (Valente et al. 2014).

Joubert Syndrome (JBTS) is a ciliopathy characterised by a midbrain and hindbrain malformation known as the “Molar Tooth Sign” (MTS). This is caused by thickened and misorientated superior cerebellar peduncles (SCPs), cerebellar vermis hypoplasia and a deepened interpeduncular fossa. Patients with JBTS can also present with additional neurological abnormalities, including further axonal tract anomalies (hypoplastic or absent corticospinal tract decussation, dysgenesis of the corpus callosum), ventriculomegaly or seizures (Bachmann-Gagescu et al. 2020; Bachmann-Gagescu et al. 2012; Bachmann-Gagescu et al. 2015; Parisi and Glass 1993; Poretti et al. 2011). Determining the pathomechanism underlying these neurological abnormalities remains a challenge due to the prominent genetic heterogeneity seen in JBTS, with over 40 associated genes. The encoded proteins all localise to distinct subcompartments of and thus likely have varying functions within the primary cilium (Bachmann-Gagescu et al. 2020; Parisi 2019). The majority, such as CC2D2A and CEP290, localise to the ciliary transition zone, a specialised domain controlling ciliary protein content (Szymanska and Johnson 2012). Other JBTS proteins localise to the basal body (TALPID3/KIAA0586 or TOGARAM1) or the ciliary membrane (INPP5E) (Latour et al. 2020; Zhang et al. 2022; Stephen et al. 2015). It can be hypothesised that dysfunction of these different proteins would contribute to a shared downstream mechanism resulting in the highly specific MTS seen in JBTS patients, as opposed to other ciliary proteins whose dysfunction does not cause JBTS. However, no such shared downstream mechanism has yet been identified.

Previous work using various models of JBTS-causative genes has shed some light on the consequences of JBTS protein dysfunction in the CNS. The primary cilium is essential for Sonic Hedgehog (Shh) signal transduction in vertebrates and many studies have identified links between JBTS protein dysfunction and Shh signalling dysregulation during early neurodevelopment. Shh is required to specify ventral cell fates during neural tube patterning, and mouse mutants for *Ift172*, *Talpid3*, *Rpgrip1l* and *Arl13b* display abnormal spinal cord patterning (Huangfu et al. 2003; Bangs et al. 2011; Vierkotten et al. 2007; Caspary et al. 2007). *Cc2d2a* mouse mutants display holoprosencephaly, a forebrain patterning defect resulting from dysregulated Shh signalling, and *Rpgrip1l* mouse mutants display forebrain developmental defects and corpus callosum agenesis (Dubourg et al. 2007; Garcia-Gonzalo et al. 2011; Besse et al. 2011; Andreu-Cervera et al. 2019). Shh is also required for the proliferation of granule cell precursors in the developing cerebellum, and mouse mutants for *Rpgrip1l* or *Talpid3* display reduced granule cell precursor proliferation leading to cerebellar vermis hypoplasia (Spassky et al. 2008; Bashford and Subramanian 2019; Wechsler-Reya and Scott 1999; Dahmane and Ruiz i Altaba 1999). Furthermore, Shh acts as an axonal guidance cue, and mouse mutants for both the Shh signalling protein *Smo* and *Arl13b* display similar aberrant SCP axonal projections (Suciu et al. 2021; Charron et al. 2003; Bourikas et al. 2005; Wilson and Stoeckli 2013).

Despite accumulating evidence that dysregulated Shh signalling as a result of JBTS protein dysfunction leads to CNS phenotypes, several studies point to additional non-Shh signalling related mechanisms as the source of these phenotypes. Aberrant development of SCPs, corticospinal tracts and the corpus callosum in *Arl13b* and *Inpp5e* mouse mutants has been linked to a dysregulation of ciliary PI3 kinase and AKT signalling (Guo et al. 2019). The *Ahi1* mouse mutant, which displays a similar though milder cerebellar hypoplasia compared to other JBTS mouse mutants and an additional cerebellar midline fusion defect, exhibits normal Shh signalling but decreased Wnt signalling at the cerebellar midline (Lancaster et al. 2011). Arl13b loss in the zebrafish results in dysregulated cerebellar Wnt signalling and an abnormal cerebellar morphology (Zhu et al. 2020). Adding another layer of complexity, the vertebrate primary cilium has been implicated in transducing a number of additional signalling pathways that have diverse roles during neurodevelopment, such as Notch and mTOR (Valente et al. 2014)

While these studies provide insight into various signalling pathways that can be disrupted upon JBTS gene dysfunction, a comprehensive analysis of the consequences of JBTS gene dysfunction during neurodevelopment is still lacking. Zebrafish are an excellent model for this purpose. There are currently multiple zebrafish mutants for JBTS-causative genes available which display various JBTS-related phenotypes, including retinal dystrophy, cystic kidneys and spinal curvature (Rusterholz et al. 2022). This is also illustrated by a recent report of 12 new zebrafish mutants affecting ciliary transition zone genes, including 8 JBTS-associated genes (Wang et al. 2022). However, a brain phenotype in these mutants has not yet been described. The zebrafish brain shares many similarities with the human brain, including a highly conserved circuitry and development of the cerebellum (Bae et al. 2009).

Hypothesising that mutations in different JBTS-causative genes would result in a shared downstream mechanism, we analysed zebrafish mutants for *cc2d2a^uw38^, cep290^fh297^, inpp5e^zh506^, talpid3^i264^* and *togaram1^zh510^*. RNA sequencing analysis revealed no signalling pathways that were similarly dysregulated in all mutants. However, all mutants displayed an enrichment of gene sets associated with CNS development and function. Despite various defects in cilia on cerebellar neurons and throughout the brain in JBTS mutants, we observed no morphological abnormalities in any brain region. Nevertheless, behavioural assays showed reduced locomotion, loss of postural control and abnormal touch-evoked escape responses.

Together with the transcriptomic results, this suggests that JBTS gene function is not essential for brain morphogenesis in zebrafish but may influence neuronal activity. While the lack of morphological abnormalities in zebrafish JBTS mutants is surprising in light of data generated in JBTS mouse mutants, it offers the unique opportunity to investigate the role of primary cilia in neuronal function beyond early developmental patterning, proliferation and differentiation.

## Results

### In search of a common transcriptomic denominator for JBTS-associated genes

To identify shared downstream consequences of JBTS gene dysfunction on the transcriptome of zebrafish larvae, we performed RNA sequencing on zebrafish mutants for the JBTS-associated genes *cc2d2a^uw38^*, *cep290^fh297^*, *togaram1^zh510^* and *talpid3^i264^* (Ben et al. 2011; Owens et al. 2008; Latour et al. 2020; Masek et al. 2022; Lessieur et al. 2019) and *inpp5e^zh506^*. These mutants were selected because they affect different sub-compartments of the primary cilium (Fig. 1A). Inpp5e localises to the ciliary membrane, Cc2d2a and Cep290 localise to the transition zone, Togaram1 localises to the base of the primary cilium and/or the ciliary tip and Talpid3 localises to the basal body (Jacoby et al. 2009; Das et al. 2015; Ben et al. 2011; Valente et al. 2006; Bachmann-Gagescu et al. 2011; Latour et al. 2020; Louka et al. 2018). Since dysfunction of all these genes causes the same highly specific CNS malformation (the MTS), our rationale was that their dysfunction must converge on a shared molecular pathway despite their evident distinct primary functions. We also analysed a zebrafish mutant for *bbs1*, which is causative for a distinct ciliopathy (Bardet-Biedl syndrome, BBS), to compare JBTS gene-specific phenotypes to another ciliopathy. Importantly, patients with BBS have intellectual disability but no CNS malformation (Forsythe et al. 2018). Generation and characterisation of the *cc2d2a^uw38^*, *cep290^fh297^*, *togaram1^zh510^, talpid3^i264^* and *bbs1^k742^* mutants have been described previously (Owens et al. 2008; Ben et al. 2011; Latour et al. 2020; Masek et al. 2022; Lessieur et al. 2019; Bachmann-Gagescu et al. 2011; Ojeda Naharros et al. 2018; Ojeda Naharros et al. 2017; Forsythe et al. 2018). *inpp5e^zh506^*mutants were generated by CRISPR editing (see Fig. S1). We used maternal zygotic (mz) mutants whenever possible (with the exception of *talpid3^i264^*, which are not viable), to prevent potential rescue of phenotypes by maternal contribution of mRNA and/or protein. Phenotypes previously described in these zebrafish mutants include curved body axis as larvae and scoliosis as adults (all mutants), retinal dystrophy (*cc2d2a^uw38^, talpid3^i264^, bbs1^k742^*) and renal cysts (*cc2d2a^uw38^, talpid3^i264^, togaram1^zh510^*) (Ben et al. 2011; Lessieur et al. 2019; Gorden et al. 2008; Owens et al. 2008; Ojeda Naharros et al. 2017; Ojeda Naharros et al. 2018; Masek et al. 2022; Latour et al. 2020). The newly generated *inpp5e^zh506^* mutant displays a similar combination of phenotypes (Fig. S1). Intriguingly, no CNS phenotype has yet been described in these mutants. In fact, with the exception of the *arl13b* mutant, CNS phenotypes have generally been little mentioned in zebrafish mutants for ciliary genes (Zhu et al. 2020; Rusterholz et al. 2022).

**Fig. 1.**
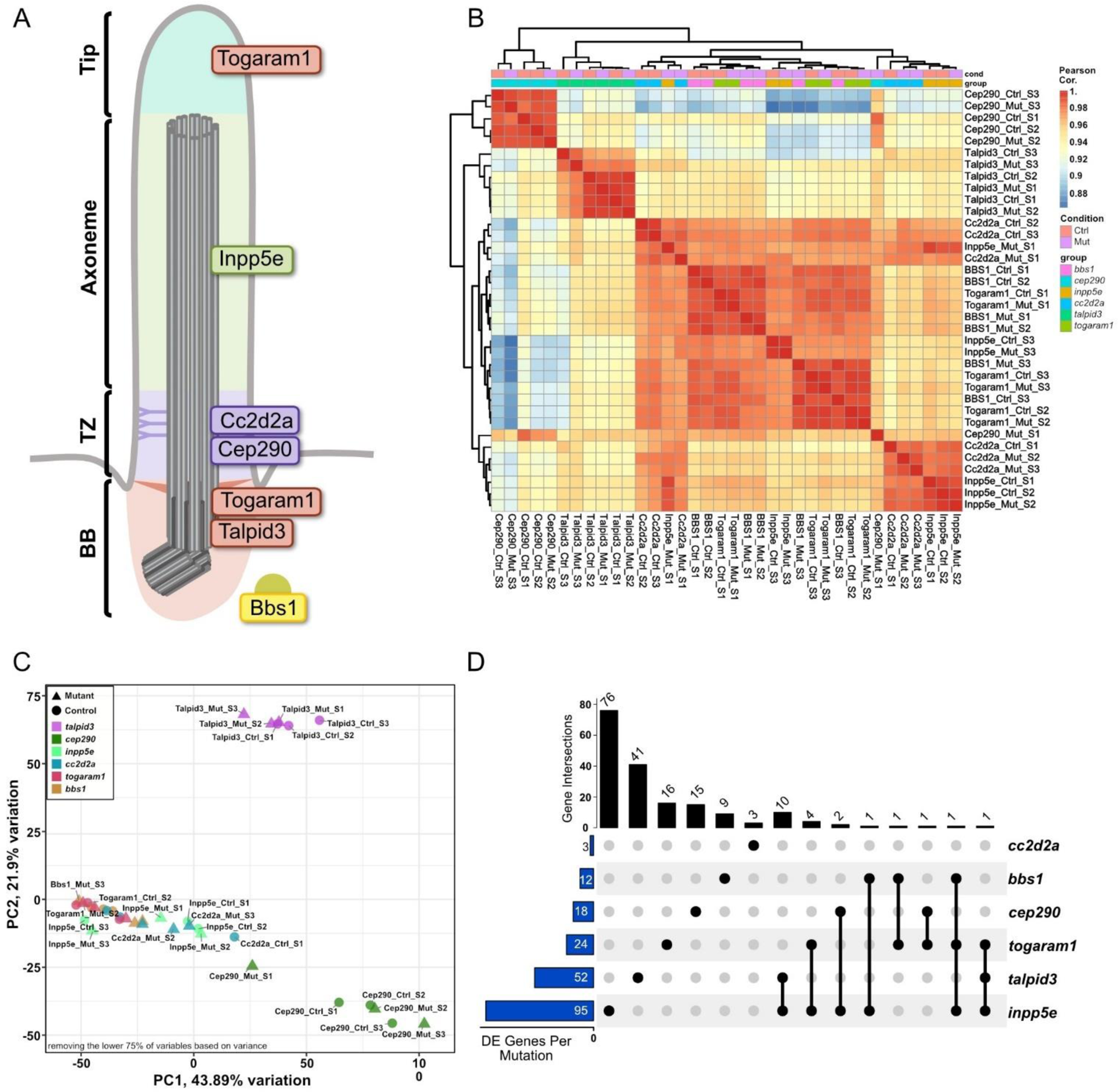
Transcriptomic comparison of zebrafish JBTS mutants. **(A)** Schematic of a cilium, outlining ciliary subcompartments (basal body (BB), transition zone (TZ), axoneme and tip). The localisation of the proteins encoded by the genes studied here is indicated. Specifically, Togaram1 and Talpid3 are localised to the BB, with Togaram1 additionally translocating to the ciliary tip, Inpp5e is localised along the ciliary membrane and Cc2d2a and Cep290 are found at the TZ. Bbs1, not associated with JBTS, functions as part of the octameric protein complex BBSome. Whole larval bulk RNA sequencing at 3 dpf was performed on zebrafish mutants harbouring mutations in one of the depicted genes. **(B)** Heat map of the Pearson correlation coefficients derived from regularised log-transformed gene counts, illustrating the overall high similarity between the different samples (correlation coefficients all above 0.88). The hierarchical clustering using average Euclidean distance reveals a pronounced batch effect in both sample pairs and zebrafish lines. **(C)** Principal component analysis focusing on the top 25% of the most variable genes, confirming a partial clustering effect attributed to the specific zebrafish lines. **(D)** UpSet plot providing a visual representation of unique and shared genes that showed significant differential expression (adjusted P-value < 0.05) in the paired differential expression analysis. The blue bars on the left show the total number of genes differentially expressed per mutant/control pair. The black bar plot on top indicates the number of genes that are commonly differentially expressed in the mutants indicated with the black dot below. Detailed information on individual genes and the intercept can be found in Tables S1 and S2.

To further characterise the selected mutants, we compared transcriptional changes by performing RNA sequencing on whole larvae at 3 days post-fertilisation (dpf). At this stage, larvae have completed most of their morphogenesis (Kimmel et al. 1995) and various cilia-dependent pathways should be active. Larvae from the same clutch (heterozygous siblings) were used as controls for each mutant sample (see methods). We observed a strong clutch-dependent effect, where samples from siblings clustered together independently of mutation status (Fig. 1B). In the principal component analysis, we found that the major source of variability in the first two components is not the mutation status but the genetic background (Fig. 1C). This finding is in agreement with zebrafish being largely outbred models and allele-specific gene expression effects due to different backgrounds have been reported (White et al. 2022). To account for these effects, we performed a paired differential expression (DE) analysis on each line individually and compared the DE genes in each mutant-control pair and between mutants. As we analysed whole larvae and were looking for systemically differentially expressed genes, only few genes reached significance (Padj. < 0.05) in this analysis and the overlap between the different mutants was even smaller (Fig. 1D, Table S1 and Table S2). Notably, the clustering for *bbs1^k742^* was indistinguishable from JBTS samples.

As Hedgehog (Hh) signalling is highly dependent on cilia, we assessed the expression levels of key Hh pathway members using the normalised pseudocounts from the RNA sequencing analysis. No systemic over/under activation of any Hh pathway members was found in any of the mutants (Fig. 2A). Since transcriptomic alterations may be tissue-specific, their effect may be diluted in this bulk assay, affecting the apparent fold change and significance levels of any potentially differentially expressed genes. Therefore, we next turned to enrichment analyses to find gene ontology terms that were enriched.

**Fig. 2.**
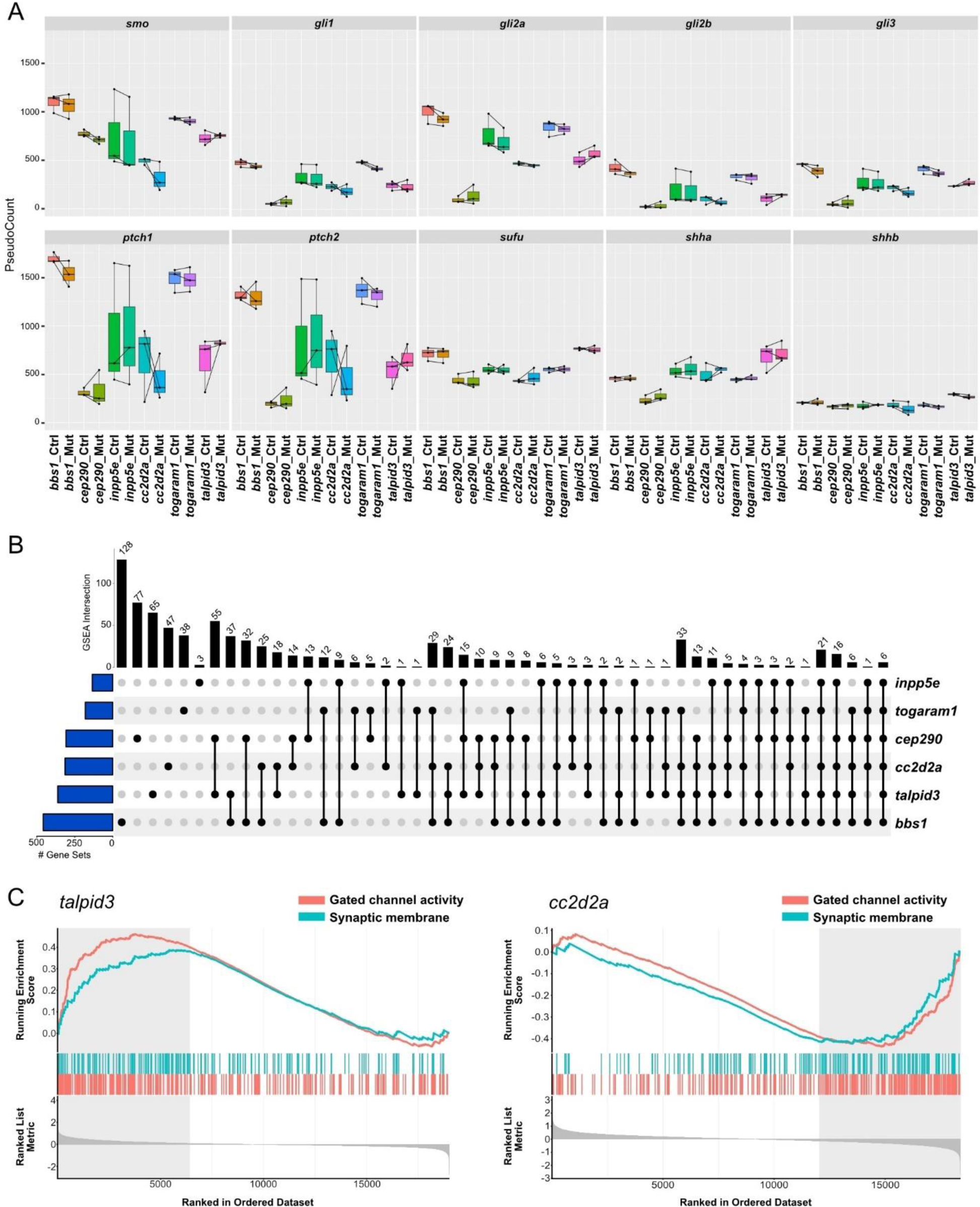
Enrichment of genes involved in central nervous system development and function in all JBTS mutants. **(A)** Box plots showing the normalised expression levels of key actors of the Shh pathway in the different mutant lines and their respective controls in the whole larval tissue analysis. Note the lack of a strong effect in any of the components in any of the mutant-sibling pairs. **(B)** Gene set enrichment analysis highlighting substantial commonality between the mutant lines. The blue bars on the left indicate the number of gene ontology terms per mutant/control pair while the black bar on top indicates the number of shared terms found in several samples indicated by the black dots. Note the terms shared between all samples (right side of the graph), which include the terms ‘gated channel activity’ and ‘synaptic membrane’. **(C)** Barcode plot detailing the position of individual genes associated to ‘gated channel activity’ (red) and ‘synaptic membrane’ (blue) along the fold change ranked list in grey at the bottom of the graph. Note that the ontology terms ‘gated channel activity’ and ‘synaptic membrane’ for *cc2d2a* and *talpid3* mutants are significantly enriched on the opposite ends of the ranked gene lists. Both terms are enriched in the up-regulated proportion in *talpid3* mutants but in the down-regulated proportion in *cc2d2a* mutants (indicated by grey boxed regions).

### JBTS mutants show an enrichment of genes involved in CNS function

We conducted enrichment analyses searching for enriched ontology terms in the different categories (Biological Process BP, Molecular Function MF and Cellular Component CC). In the overrepresentation analysis (ORA) with threshold P-value < 0.01 and absolute FoldChange > 1.5, we found very few genes that were significantly enriched in more than one mutant (Fig. S2 and Table S3). Several mutants showed an enrichment of terms associated with vision (“visual perception” or “detection of visible light”). This is consistent with the previously described retinal phenotype of *cc2d2a^uw38^* and *talpid3^i264^*mutants at 4 and 5 dpf. Our results indicate that transcriptional changes are already present earlier at 3 dpf, even if the signal transmission of photoreceptors starts only later around 84 hours post-fertilisation (hpf) (Biehlmaier et al. 2003).

Next, we evaluated ontology terms that were found to be enriched using the fold change ranked gene list to assess whether gene sets were non-randomly distributed at one extreme of the list (GSEA). Strikingly, we found in the GSEA that the MF ontology term ‘gated channel activity’ and the CC ontology term ‘synaptic membrane’ were enriched in all mutants (Fig. 2B and Table S4). It is noteworthy that the enrichment of the terms is not in the same direction in all mutants, as in *cc2d2a^uw38^*, *bbs1^k742^*and *togaram1^zh510^* the genes associated with these terms were in the downregulated proportion, whereas in *talpid3^i264^*, *inpp5e^zh506^* and *cep290^fh297^* they were found in the upregulated proportion (Fig. 2C and Table S4). Additionally, terms associated with axon guidance were found to be shared by most of the mutants (Table S4). These findings suggest a defect in CNS development and/or function in all mutants. We therefore next aimed to characterise the brain phenotype of JBTS zebrafish mutants.

### Cerebellar development is unaffected in JBTS mutants despite abnormal primary cilia

As the defining feature of JBTS is a highly specific midbrain-hindbrain malformation (the MTS), we focused first on analysing the cerebellum of zebrafish JBTS mutants. The zebrafish cerebellum is organised into three distinct cell layers, the ventral granule cell layer (GCL), Purkinje cell layer (PCL) and dorsal molecular layer (ML), similar to what is found in other vertebrates, including mouse and human. Within the GCL, granule cells receive excitatory inputs from mossy fibres. The granule cell axons, named parallel fibres, extend dorsally to the ML where they contact the intricately branched dendritic trees of Purkinje cells. Purkinje cell dendrites also receive excitatory inputs from climbing fibres. Information from mossy fibres and climbing fibres is integrated by Purkinje cells, whose soma reside in the PCL, and the information from Purkinje cells is conveyed to the brain via eurydendroid cells residing ventrally to Purkinje cells. Eurydendroid cells are equivalent to mammalian deep cerebellar nuclei (DCN). The zebrafish cerebellum has at least two subsets of eurydendroid cells, a large subset expressing *oligodendrocyte transcription factor 2 (olig2)*, and a smaller subset expressing *calretinin* (Fig. 3A-B) (Bae et al. 2009; Biechl et al. 2016; McFarland et al. 2008).

**Fig. 3.**
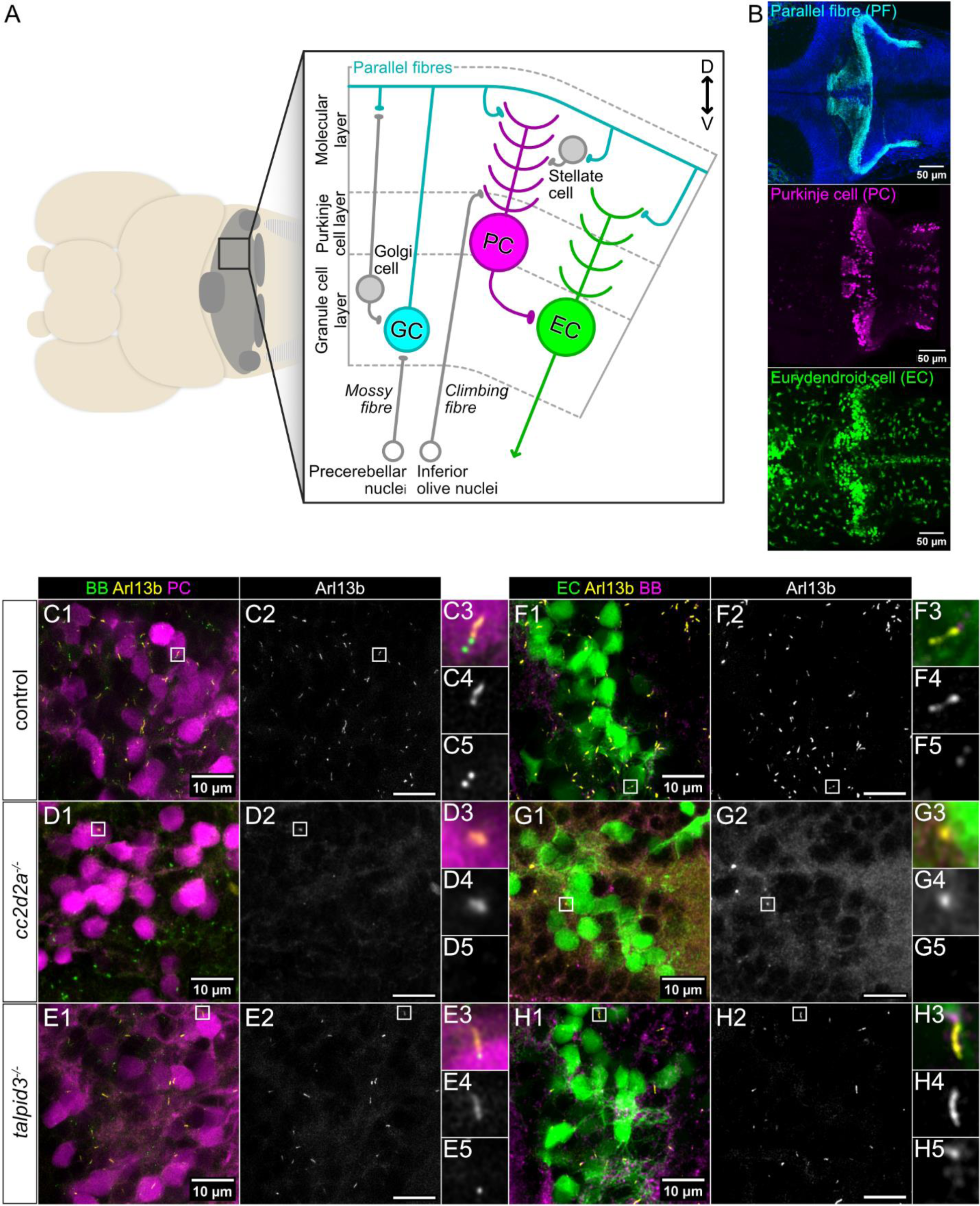
Loss of primary cilia in cerebellar neurons of *cc2d2a* and *talpid3* mutants. **(A)** Schematic representation of cerebellar circuits in a zebrafish larva at 5 dpf. **(B)** Representative whole-mount maximum projection confocal immunofluorescence images showing parallel fibres of granule cells (anti-Vglut1 – cyan), Purkinje cells (anti-Pvalb7 – magenta) and *olig2+* eurydendroid cells (*Tg(olig2:EGFP)* transgenic line – green) in the cerebellum of wild-type zebrafish larvae at 5 dpf. Images show a dorsal view with anterior to the left. **(C-H)** Whole-mount maximum projection confocal immunofluorescence images showing primary cilia labelled with the *Tg(ubi:Arl13b-mCherry)* transgenic line (Arl13b – yellow) and basal bodies (BB – green) labelled using anti-γ-Tubulin (D, F-H) or using the *Tg(β-actin:GFP-centrin)* transgenic line (C, E). Purkinje cells (PC – magenta) are marked using anti-Pvalb7 (C-E) and eurydendroid cells (EC – green) are marked by the *Tg(olig2:EGFP)* transgenic line (F-H). Note the strong decrease in cilia numbers in both mutants (*cc2d2a^−/−^* in D and G; *talpid3^−/−^*in E and H) compared to control (C and F). Insets show a magnified view of one representative cilium (from top to bottom: composite image (C3-H3), Arl13b (C4-H4) and BB (C5-H5)). Images show a dorsal view of 5 dpf larvae with anterior to the left. Scale bars are 10 µm.

We chose to concentrate on two mutants for the JBTS genes *cc2d2a* and *talpid3*, as these genes encode proteins localising to different compartments of and with different functions within the primary cilium (Ojeda Naharros et al. 2018; Ojeda Naharros et al. 2017). Moreover, these mutants displayed opposite effects on the transcriptome with downregulation of CNS-associated terms in *cc2d2a* but upregulation in *talpid3*.

We first analysed primary cilia in the cerebellum of *cc2d2a^−/−^* and *talpid3^−/−^*at 5 dpf, as cerebellar neuron differentiation begins at 3 dpf and layer formation is evident at 5 dpf (Bae et al. 2009; Biechl et al. 2016). We crossed *cc2d2a* and *talpid3* lines with a *Tg(ubi:Arl13b-mCherry)^zh406^*transgenic line that we generated (see methods), which ubiquitously expresses an Arl13b-mCherry fusion protein. Arl13b is a ciliary-enriched GTPase localised along the ciliary axoneme and can therefore be used to label cilia. To identify the different cerebellar neuronal subtypes, we used antibodies against Pvalb7 to label Purkinje cells and the *Tg(olig2:EGFP)* transgenic fluorescent reporter line to label *olig2+* eurydendroid cells (McFarland et al. 2008; Shin et al. 2003). Whole-mount immunofluorescence imaging indicated that *cc2d2a^−/−^* and *talpid3^−/−^* have fewer Arl13b-positive primary cilia on Purkinje and eurydendroid cells compared to controls (Fig. 3C-H). This result was confirmed using an antibody against endogenous Arl13b (Fig. S3A-D).

We next investigated whether the reduction in Arl13b-positive primary cilia in *cc2d2a^−/−^* and *talpid3^−/−^* resulted in changes to cerebellar morphology. We crossed our mutant lines with the *Tg(tagRFP-T:PC:GCaMP5G)* fluorescent reporter line, in which tagRFP-T is expressed in Purkinje cells (Matsui et al. 2014) and found that the morphology of the Purkinje cell layer in *cc2d2a^−/−^* and *talpid3^−/−^* is unaffected (Fig. 4A-C). Furthermore, the number of Purkinje cells per cerebellar hemisphere is not significantly changed in either mutant compared to controls (mean cell number in control = 126.8±23.13 cells, *cc2d2a^−/−^* = 125.8±16.86 cells, *talpid3^−/−^* = 130.5±18.49 cells) (Fig. 4D and Fig. S4). To assess parallel fibre-Purkinje cell synapses, we used an antibody against the vesicular glutamate transporter 1 (Vglut1). The distribution and density of Vglut1-positive parallel fibre synapses in the ML was unchanged in both mutants (Fig. 4E-G), as there was no difference in the normalised Vglut1-positive fluorescence area in *cc2d2a^−/−^* and *talpid3^−/−^* compared to controls (mean normalised area in controls = 0.0222±0.0032 µm, *cc2d2a^−/−^* = 0.0222±0.0026 µm, *talpid3^−/−^*= 0.0206± 0.0022 µm) (Fig. 4H).

**Fig. 4.**
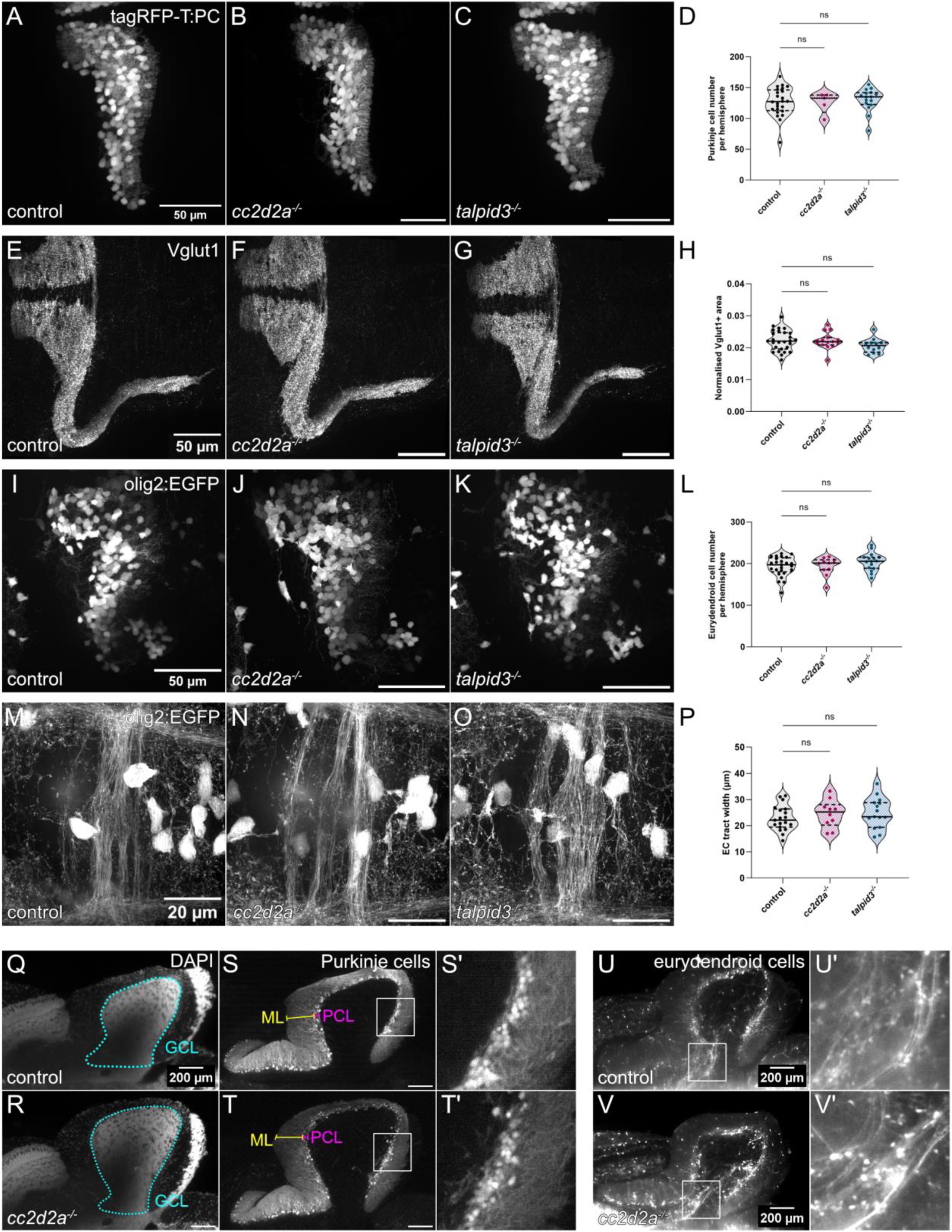
Morphological development of the cerebellum is unaffected in *cc2d2a* and *talpid3* mutants. **(A-C)** Whole-mount maximum projection confocal images showing tagRFP-T-positive Purkinje cells in control (A), *cc2d2a^−/−^*(B) and *talpid3^−/−^* (C) *Tg(tagRFP-T:PC:GCaMP5G)* larvae at 5 dpf. Note the similar morphology of the PCL in mutants and controls. **(D)** Violin plot showing no statistically significant difference in the number of tagRFP-T-positive Purkinje cells per cerebellar hemisphere in mutants compared to controls. Control n = 24 larvae, *cc2d2a^−/−^*n = 5 larvae (P = 0.99), *talpid3^−/−^* n = 16 larvae (P = 0.82). **(E-G)** Whole-mount maximum projection confocal immunofluorescence images showing normal morphology of Vglut1-positive parallel fibres in control (E), *cc2d2a^−/−^* (F) and *talpid3^−/−^* (G) larvae at 5 dpf. **(H)** Violin plot showing no statistically significant difference in the normalised Vglut1-positive fluorescence area in mutants compared to controls. Control n = 27 larvae, *cc2d2a^−/−^* n = 15 larvae (P = 1.00), *talpid3^−/−^* n = 12 larvae (P = 0.21). **(I-K)** Whole-mount maximum projection confocal images showing EGFP-positive eurydendroid cells in control (I), *cc2d2a^−/−^* (J) and *talpid3^−/−^* (K) *Tg(olig2:EGFP)* larvae at 5 dpf. The morphology of the eurydendroid cell cluster is similar in mutants and controls. **(L)** Violin plot showing no statistically significant difference in the number of EGFP-positive eurydendroid cells per cerebellar hemisphere in mutants compared to controls. Control n = 25 larvae, *cc2d2a^−/−^* n = 11 larvae (P *=* 0.99), *talpid3^−/−^*n = 16 larvae (P = 0.20)**. (M–O)** Whole-mount maximum projection confocal images showing EGFP-positive eurydendroid cell axons in control (M), *cc2d2a^−/−^* (N) and *talpid3^−/−^* (O) *Tg(olig2:EGFP)* larvae at 5 dpf. **(P)** Violin plot showing quantification of the EGFP-positive axon tract thickness, where no statistically significant difference is observed in mutants compared to controls. Control n = 19 larvae, *cc2d2a^−/−^* n = 10 larvae (P = 0.65), *talpid3^−/−^* n = 15 larvae (P = 0.50). For violin plots in D, H, L and P, each data point represents one larva. Violin plots represent median (thick line) and quartiles (dashed line). ns = not significant. Ordinary one-way ANOVA with post-hoc Dunnett’s multiple comparisons test. For A-O, images show a dorsal view of 5 dpf larvae with anterior to the left. For A-K, image scale bars are 50 µm and for M-O, image scale bars are 20 µm. **(Q-T)** Representative maximum projection mesoSPIM images showing an optical sagittal section of DAPI-stained nuclei and tagRFP-T-positive Purkinje cells in the cerebellum of *Tg(tagRFP-T:PC:GCaMP5G)* 11-12 wpf control (Q, S) and *cc2d2a^−/−^* (R, T) fish. S’ and T’ show higher magnification images of boxed regions in S and T, respectively. The GCL is outlined with a cyan line, the PCL is indicated with a magenta bracket and the ML with a yellow bracket. The morphology of all layers is unchanged in mutants. **(U-V)** Maximum projection mesoSPIM images showing an optical sagittal section of olig2:EGFP-positive eurydendroid cells in the cerebellum of *Tg(olig2:EGFP)* 11-12 wpf control (U) and *cc2d2a^−/−^* (V) fish. Eurydendroid cells and axons appear normal in mutants compared to controls. U’ and V’ show higher magnification images of boxed regions in U and V, respectively. For Q-V, images show a sagittal view with anterior to the left and scale bars are 200 µm.

We next crossed the *Tg(olig2:EGFP)* transgene into our mutants to analyse *olig2+* eurydendroid cells. We found the morphology and the number of *olig2+* eurydendroid cells per cerebellar hemisphere to be unaffected in *cc2d2a^−/−^* and *talpid3^−/−^* compared to controls (mean cell number in control = 192.6±23.52 cells, *cc2d2a^−/−^*= 193.7±21.66 cells, *talpid3^−/−^* = 204.3±21.11 cells) (Fig. 4I-L and Fig. S4). A hallmark of JBTS is the abnormal morphology of SCPs, which represent the cerebellar output from the DCN in mammals. As eurydendroid cells are equivalent to mammalian DCN (Bae et al. 2009), we therefore analysed eurydendroid cell axons leaving the cerebellum anteriorly and crossing the midline using the same transgenic *Tg(olig2:EGFP)* line. This eurydendroid cell axonal tract revealed variable morphologies within control, *cc2d2a^−/−^* and *talpid3^−/−^* groups, however no consistent morphological abnormalities or changes in tract width could be seen in *cc2d2a^−/−^* and *talpid3^−/−^* compared to control (mean tract width in control = 22.90±4.82µm, *cc2d2a^−/−^*= 24.55±5.33µm, *talpid3^−/−^* = 24.79±5.81µm) (Fig. 4M-P and Fig. S3E-H).

Seeing as we identified no differences in cerebellar morphology between control and mutant larvae, we next questioned whether cerebellar abnormalities may not appear until later developmental stages. As *talpid3^−/−^*are not viable, we analysed cerebellar morphology of zygotic *cc2d2a* juveniles at 11-12 weeks post-fertilisation (wpf) using lightsheet mesoSPIM imaging of whole cleared brains (Voigt et al. 2019; Vladimirov et al. 2023 preprint). At this later stage, we also observed no abnormalities in the overall morphology of the cerebellum or in the organisation of the PCL in *cc2d2a^−/−^*(Fig. 4Q-T, Movies 1 and 2). Likewise, organisation of eurydendroid cells and their axons, highlighted by the *Tg(olig2:EGFP)* transgene, appeared normal in *cc2d2a^−/−^*(Fig. 4U-V).

Given the unexpected lack of a cerebellar morphological defect in *cc2d2a^−/−^* and *talpid3^−/−^* despite abnormal primary cilia, we further analysed cerebellar morphology in the remaining mutants (*inpp5e^−/−^, cep290^−/−^, togaram1^−/−^* and *bbs1^−/−^*) using antibodies against Pvalb7 to label Purkinje cells, Calretinin to label eurydendroid cells and Vglut1 to label parallel fibres. As for *cc2d2a^−/−^* and *talpid3^−/−^*, we observed no anomalies in cerebellar morphology in any of the other mutants (Fig. S5). Taken together, these results indicate that loss of JBTS gene function does not affect cerebellar development in zebrafish despite abnormalities in primary cilia.

### JBTS mutants have normal brain morphology despite abnormalities in CNS motile and primary cilia

Given the observed ciliary defects and transcriptomic changes, we expanded our analysis to the entire brain of the zebrafish JBTS mutants. We first determined whether cilia, both primary and motile, were affected in the brain of *cc2d2a^−/−^* and *talpid3^−/−^*.

Whole-mount immunofluorescence imaging shows that controls have abundant primary cilia in the forebrain and midbrain parenchyma. As in the cerebellum, *cc2d2a^−/−^* have fewer Arl13b-positive primary cilia in the forebrain and midbrain parenchyma compared to controls. Interestingly, primary cilia appear to be only moderately reduced in the forebrain parenchyma of *talpid3^−/−^* but strongly reduced in the midbrain parenchyma (Fig. 5B-G).

**Fig. 5.**
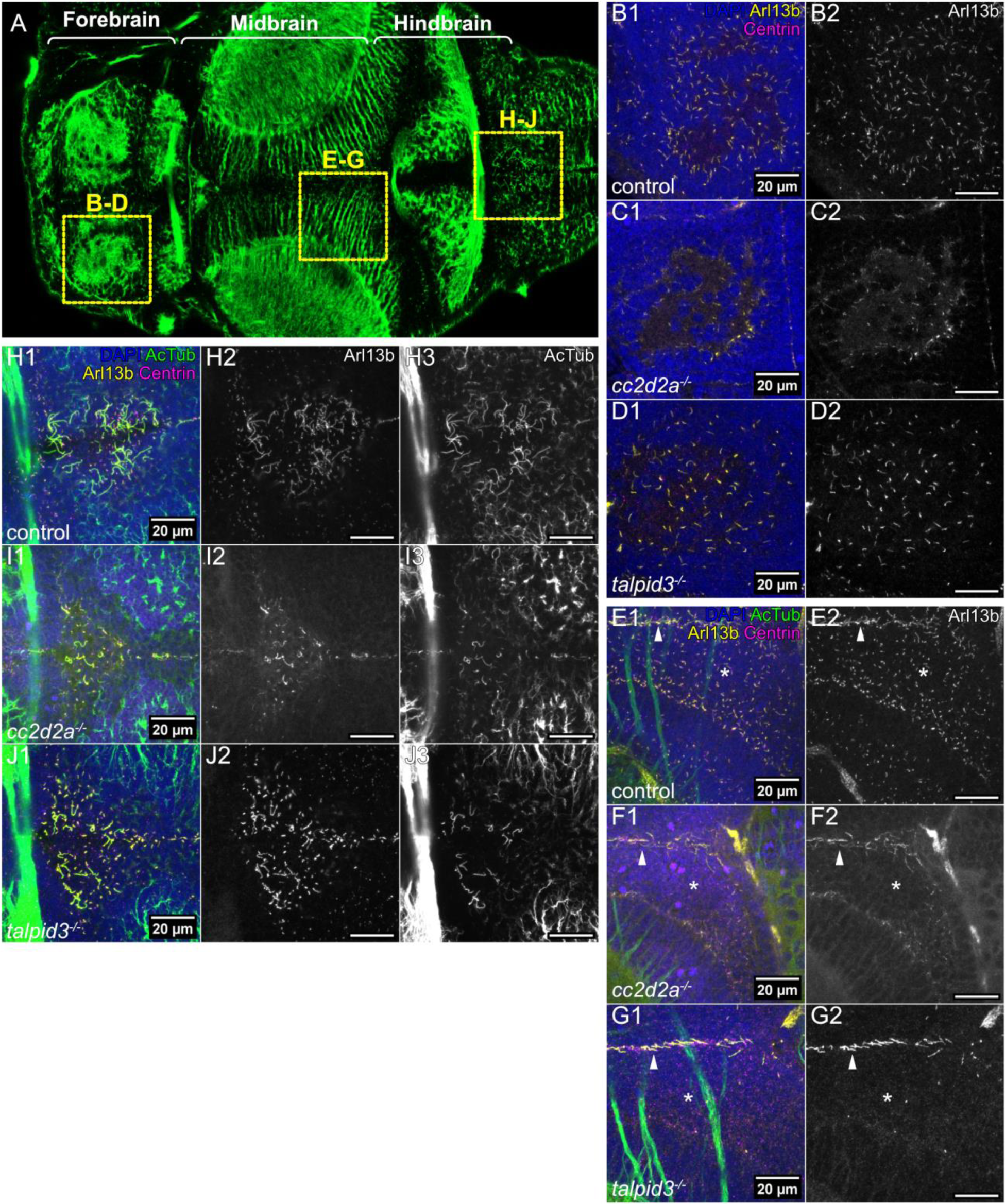
Abnormal cilia in the brain of *cc2d2a* and *talpid3* mutants. **(A)** Whole-mount confocal image of a 5 dpf zebrafish larva immunostained with anti-acetylated tubulin, providing orientation for the figure. Anterior is to the left and boxes show the analysed brain regions in the subsequent panels. **(B-G)** Whole-mount maximum projection confocal immunofluorescence images of the forebrain (B-D) and midbrain (E-G), showing **primary** cilia labelled with anti-Arl13b (Arl13b – yellow) and basal bodies labelled with anti-Centrin (Centrin – magenta) in control, *cc2d2a^−/−^* and *talpid3^−/−^* larvae at 5 dpf. In E-G, Arl13b-positive and acetylated tubulin-positive motile cilia are visible at the midline (indicated with white arrow heads), while asterisks indicate location of Arl13b-positive primary cilia in the midbrain. In both *cc2d2a^−/−^* and *talpid3^−/−^*larvae, there is a marked reduction in primary cilia. **(H-J)** Whole-mount maximum projection confocal immunofluorescence images of the hindbrain ventricle, using anti-acetylated tubulin (AcTub – green) and anti-Arl13b (Arl13b – yellow) to label **motile** cilia and anti-Centrin (Centrin – magenta) to label basal bodies. Motile cilia are reduced in both *cc2d2a^−/−^* and *talpid3^−/−^* larvae compared to controls. All images show a dorsal view of 5 dpf larvae with anterior to the left. Scale bars are 20 µm.

In the hindbrain ventricle, controls show long motile cilia that can be immunolabelled with antibodies against Arl13b and acetylated tubulin and shorter Arl13b-positive primary cilia that show fainter immunostaining for acetylated tubulin. Both *cc2d2a^−/−^* and *talpid3^−/−^* have fewer motile cilia, with fewer primary cilia visible in *cc2d2a^−/−^* than in *talpid3^−/−^* (Fig. 5H-J). Within the midbrain and forebrain ventricles, *cc2d2a^−/−^*have fewer motile cilia with reduced immunostaining for Arl13b, while *talpid3^−/−^*motile cilia appear unaffected (Fig. S6). Taken together, these results indicate that there is a global reduction in primary and motile cilia within the brain of *cc2d2a^−/−^* and *talpid3^−/−^*compared to controls, however regional differences in cilia abundance vary between *cc2d2a^−/−^* and *talpid3^−/−^* and within each mutant between different regions.

Other JBTS mutants also show variable ciliary phenotypes throughout the brain. In the forebrain, primary cilia in *inpp5e^−/−^* appear shorter while midbrain primary cilia are unaffected. In *cep290^−/−^*, we observed a slight reduction in primary cilia number in the forebrain and midbrain. Primary cilia within the midbrain and forebrain of *togaram1^−/−^* appear shorter and slightly reduced in number. In the forebrain and midbrain of *bbs1^−/−^* we observe some shorter primary cilia (Fig. S7). Motile cilia appear normal in the forebrain and midbrain ventricles in *inpp5e^−/−^*, while hindbrain ventricle motile cilia are shorter and fewer. *cep290^−/−^* appear to have slightly fewer motile cilia. In *togaram1^−/−^*, we see a substantial reduction in acetylated tubulin in all motile cilia, which are also shorter and fewer. *bbs1^−/−^*also have fewer motile cilia with reduced levels of acetylated tubulin (Fig. S8). This data supports our findings in *cc2d2a^−/−^* and *talpid3^−/−^*, indicating wide-spread but variable anomalies of primary and motile cilia throughout the brain of JBTS mutants.

Given this global alteration of cilia in the brains of JBTS mutants, we next sought to determine whether brain morphology was affected in *cc2d2a^−/−^*and *talpid3^−/−^*. We used antibodies against HuC/HuD to label neurons at 5 dpf and observed no changes in overall brain morphology in either mutant (Fig. 6A-C). To rule out different effects on subregions of the brain, we quantified forebrain, midbrain and hindbrain area normalised to whole brain area (Fig. 6E-F). This analysis revealed that the mean normalised area of these three regions is not significantly different in *cc2d2a^−/−^* compared to controls (Table S5). Similar findings were identified for *talpid3^−/−^* with the exception of a minimal but statistically significant difference in the mean normalised forebrain area (Table S6). We then used antibodies against acetylated tubulin to label axonal tracts and against synaptic vesicle 2 (SV2) to label synapses to further investigate neural circuit formation. This analysis also did not show any difference in the morphology of axonal tracts or the synaptic neuropil in *cc2d2a^−/−^*or *talpid3^−/−^* (Fig. 6G-L).

**Fig. 6.**
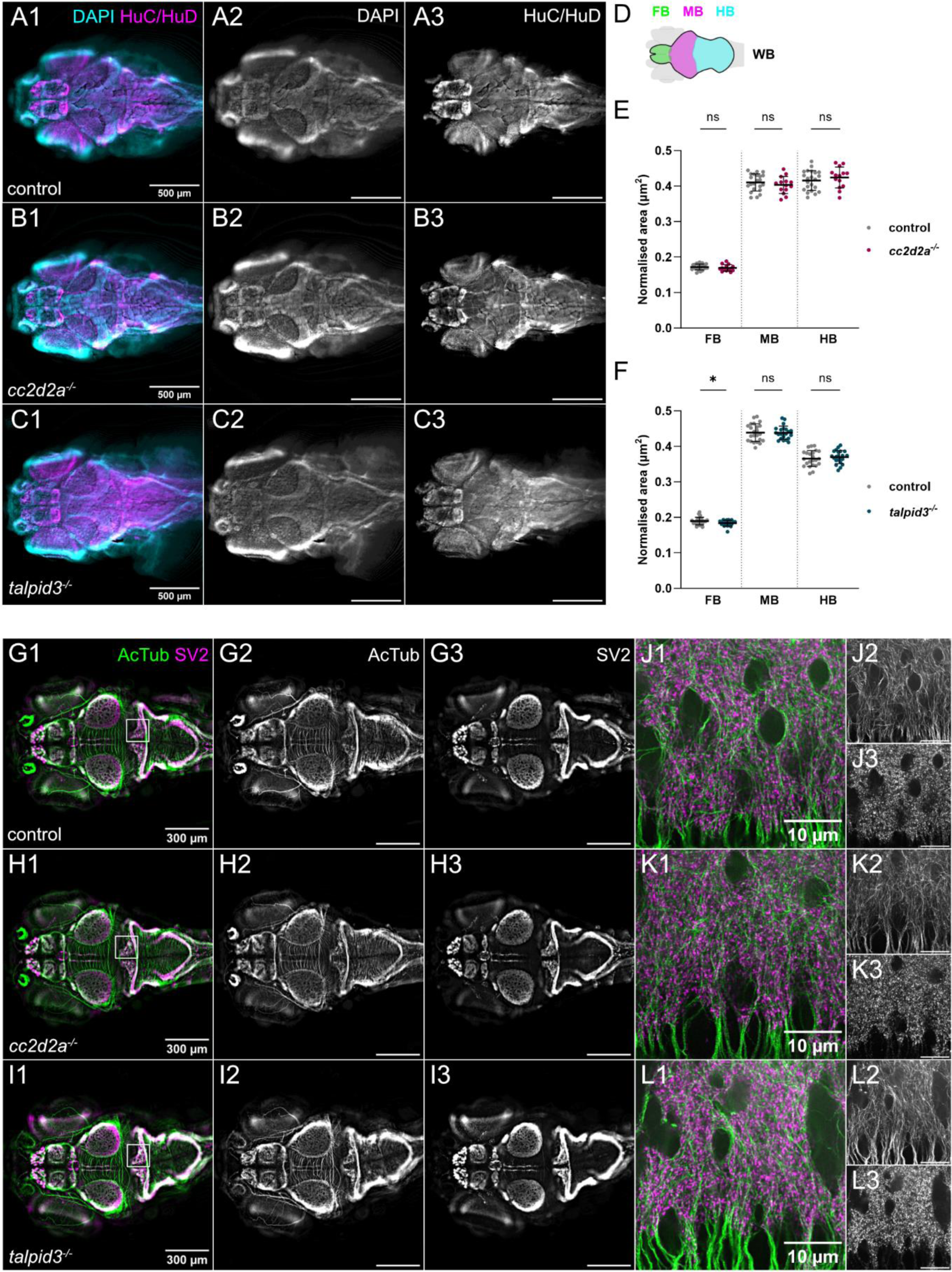
Brain morphology and size is normal in *cc2d2a* and *talpid3* mutants. **(A-C)** Whole-mount single optical slice widefield immunofluorescence images showing brain morphology of control (A), *cc2d2a^−/−^* (B) and *talpid3^−/−^* (C) larvae at 5 dpf, stained with anti-HuC/HuD to label neurons (HuC/HuD – magenta) and DAPI to counterstain nuclei (DAPI – cyan). Note that the morphology of the brain in *cc2d2a^−/−^* and *talpid3^−/−^*larvae is comparable to controls. **(D)** Schematic showing regions of interest used for brain area quantification. **(E-F)** Scatter plots showing normalised area of the forebrain, midbrain and hindbrain regions in control vs. *cc2d2a^−/−^* larvae (E) and control vs. *talpid3^−/−^* larvae (F) at 5 dpf. There is no statistically significant difference in normalised brain area in *cc2d2a^−/−^* larvae compared to controls. In *talpid3^−/−^* larvae, there is a minimal but statistically significant difference in normalised forebrain area compared to controls. Each data point represents one larva. Error bars are mean±s.d. ns, not significant; * P ≤ 0.05. Unpaired t test. For (E), control n = 22, *cc2d2a^−/−^* n = 14 larvae. For (F), control n = 23, *talpid3^−/−^* n = 22 larvae. **(G-L)** Whole-mount single optical slice widefield immunofluorescence images showing an overview of axonal tracts labelled with anti-acetylated tubulin (AcTub – green) and synaptic neuropil labelled with anti-synaptic vesicle 2 (SV2 – magenta) in control (G), *cc2d2a^−/−^* (H) and *talpid3^−/−^* (I) larvae at 5 dpf. (J-L) shows higher resolution whole-mount single optical slice confocal immunofluorescence images of boxed region in (G-I). The morphology of axonal tracts and synaptic neuropil appears unchanged in both mutants compared to controls. All images show a dorsal view of 5 dpf larvae with anterior to the left. Scale bars are 500 µm in A-C, 300 µm in G-I and 10 µm in J-L.

As before, given the surprising absence of CNS phenotype in *cc2d2a*^−/−^ and *talpid3^−/−^*, we turned to the remaining mutants in our collection and analysed brain morphology and neuropil organisation in *inpp5e^−/−^, cep290^−/−^*, *togaram1^−/−^* and *bbs1^−/−^*. We observed no changes in the gross morphology of the brain and subregions on immunostaining with anti-HuC/HuD or in the organisation of axonal tracts and synapses on immunostaining with anti-acetylated tubulin and anti-SV2, respectively (Fig. S9).

Taken together, our data indicate that brain development in zebrafish JBTS mutants occurs normally despite abnormal primary cilia in the forebrain, midbrain and hindbrain and abnormal motile cilia in the ventricles. However, we cannot rule out that higher resolution analyses of axonal tracts or synaptic neuropil may reveal subtle changes in neural circuit development.

### JBTS mutants have abnormal swimming behaviour

Our RNA sequencing analysis indicates that JBTS mutants have an enrichment of GO terms related to neuronal activity. Zebrafish larvae display a wide range of behaviours that are controlled by neural circuits in the brain and spinal cord. We therefore next analysed larval locomotion and escape responses to investigate alterations in neuronal activity in *cc2d2a^−/−^*and *talpid3^−/−^*.

We analysed locomotion at 3 dpf and 6 dpf using automatic movement tracking with the Zebrabox during three sequential five minute periods of darkness-light-darkness. At 3 dpf, control larvae showed characteristic infrequent, erratic bouts of spontaneous swimming (Buss and Drapeau 2001). We could see no significant difference in the locomotion of *cc2d2a^−/−^*or *talpid3^−/−^* compared to controls, indicated by comparable distance travelled over time and total distance travelled, despite a trend towards decreased movement in mz *cc2d2a^−/−^* (Fig. S10, Table S7 and Table S8). However, we observed a statistically significant difference in locomotion in *cc2d2a^−/−^* mutants and controls at 6 dpf. Here, control larvae displayed increased locomotion during darkness and decreased locomotion during light and their swimming behaviour followed the characteristic “beat-and-glide” swimming and thigmotaxis that are expected at 6 dpf (Movie 3) (Buss and Drapeau 2001; Basnet et al. 2019). In contrast, *cc2d2a^−/−^* had significantly reduced locomotion throughout compared to controls, and this reduction was most evident during darkness periods. To rule out that this reduced locomotion was due to the lack of a swim bladder or more severe body curvature of these mutants, we also analysed zygotic *cc2d2a^−/−^* with a swim bladder which showed only very mild body curvature. These mildly affected zygotic *cc2d2a*^−/−^ also showed a significantly reduced locomotion compared to controls (Fig. 7A-B and 7E, Table S7). In addition, both mz *cc2d2a^−/−^* and zygotic *cc2d2a^−/−^*were sometimes lying on their side at the bottom of the well, suggesting abnormalities in postural control (Movies 4 and 5). No significant differences in locomotion were observed in *talpid3^−/−^*, however a slight reduction in locomotion after the light to darkness transition could be seen (Fig. 7C-E, Table S8).

**Fig. 7.**
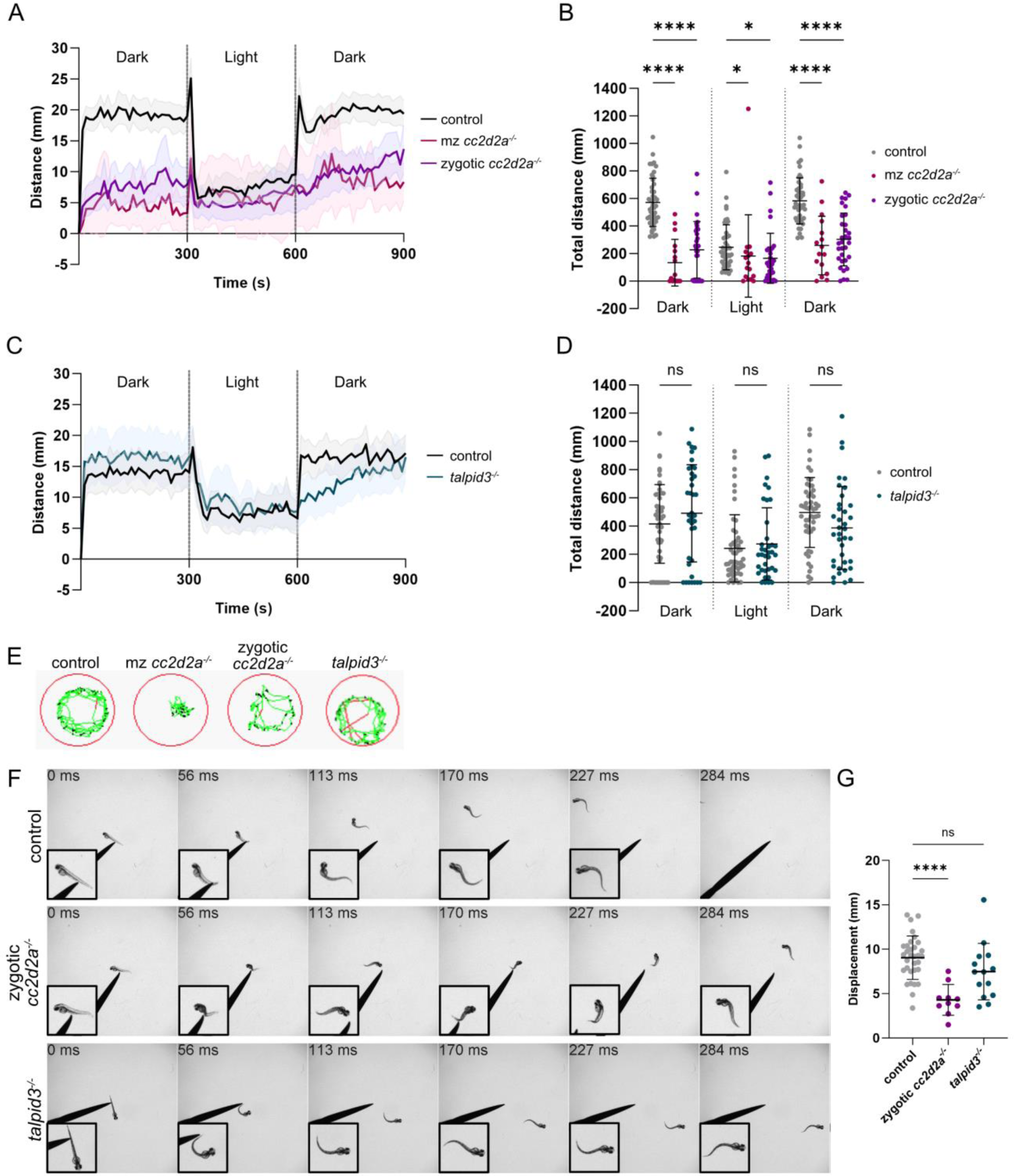
Abnormal swimming behaviour in JBTS mutants. **(A)** Plot showing the mean distance travelled during 3 sequential 5-minute periods of darkness, light, darkness in mz *cc2d2a^−/−^* and milder zygotic *cc2d2a^−/−^* larvae compared to controls. Total distance moved is recorded during 10 second intervals and this distance is plotted over time. Note that both mz *cc2d2a^−/−^* and zygotic *cc2d2a^−/−^* larvae travel less distance than controls during both dark periods. Error shown with the coloured areas represent the 95% CI. **(B)** Scatter plot showing the total distance travelled during the 5-minute periods of darkness, light and darkness in mz *cc2d2a^−/−^* and zygotic *cc2d2a^−/−^*larvae compared to controls. The total distance travelled is significantly reduced in both mz *cc2d2a^−/−^* and zygotic *cc2d2a^−/−^*. Error bars are mean±s.d. ns, not significant; * P ≤ 0.05; **** P ≤ 0.0001. Kruskal-Wallis test with post-hoc Dunn’s multiple comparisons test. Control n = 46, mz *cc2d2a^−/−^* n = 16, zygotic *cc2d2a^−/−^* n = 32 larvae. **(C)** Similar plot as in (A) but for *talpid3^−/−^*larvae compared to controls. The distance travelled throughout the experiment does not differ between mutants and controls. Error shown with the coloured areas represent the 95% CI. **(D)** Scatter plot showing the total distance travelled by *talpid3^−/−^* larvae compared to controls. The total distance is not significantly reduced in *talpid3^−/−^* larvae (trend towards decrease in the second dark phase not significant). Error bars are mean±s.d. ns, not significant. Unpaired t test. Control n = 51, *talpid3^−/−^* n = 37 larvae. **(E)** Representative 1-minute traces showing locomotion in one control, one mz *cc2d2a^−/−^,* one zygotic *cc2d2a^−/−^* and one *talpid3^−/−^* larva. Black traces represent inactivity, green traces represent normal activity and red traces represent fast activity. **(F)** Representative montages of the touch-evoked escape response (TER) in control, zygotic *cc2d2a^−/−^* and *talpid3^−/−^*larvae at 2 dpf. Insets show magnified view of the larvae. While the control and *talpid3^−/−^* larvae are able to escape the stimulus, the abnormal swimming of the zygotic *cc2d2a^−/−^* larvae (circular with partial swimming on the side or upside down) prevents it from escaping from the stimulus in the same manner. Please note the only mild curvature of the larva, which should not suffice to explain the tightly curved swimming trace observed. **(G)** Scatter plot showing displacement of control, zygotic *cc2d2a^−/−^* and *talpid3^−/−^* larvae during the TER. Zygotic *cc2d2a^−/−^* exhibit a significantly reduced displacement compared to controls. The trend towards decreased displacement in *talpid3*−/− does not reach statistical significance. Error bars are mean±s.d. not significant; **** P ≤ 0.0001. Ordinary one-way ANOVA with post-hoc Dunnett’s multiple comparisons test. Control n = 32, zygotic *cc2d2a^−/−^* n = 10 (P = <0.0001), *talpid3^−/−^* n = 14 (P = 0.1146) larvae.

We next analysed the touch-evoked escape response (TER) at 2 dpf (Fig. 7F-G), a well-studied stereotypical zebrafish behaviour initiated by a single touch to the tail. The TER in control larvae involved a fast swimming action away from the stimulus, as expected. Zygotic *cc2d2a^−/−^* exhibited abnormal TERs, swimming in a circular fashion around their original starting position and sometimes swimming on the side or upside down, which is reflected in a significantly reduced displacement. As zygotic *cc2d2a^−/−^* display only a mild curvature, these abnormal TERs cannot be fully explained by the minimal curvature of their tail tip. Conversely, *talpid3^−/−^*displayed a normal TER with no significant reduction in displacement (mean displacement in control = 9.039±2.451 mm, zygotic *cc2d2a^−/−^* = 4.284±1.733 mm, *talpid3^−/−^* = 7.472±3.183 mm).

Taken together, the presence of these abnormal behaviours despite normal brain morphology may indicate a deficit in neuronal activity in zebrafish JBTS mutants.

## Discussion

A crucial role for primary cilia during early CNS development, notably patterning and proliferation, is suggested through the study of mouse models harbouring mutations in ciliary genes. This aligns with the primary cilium’s role as a sensory organelle that transduces developmental signalling pathways like Hh or Wnt. However, their role in differentiated neurons is less clear even though primary cilia are present on differentiated neurons and astrocytes (Sterpka and Chen 2018). This limited understanding is in part due to the major malformations observed in mouse cilia mutants that obscure a subsequent role for primary cilia in differentiated neurons. Importantly, the CNS phenotypes associated with ciliary dysfunction in humans tend to be less severe than those observed in mice. Indeed, the MTS characteristic of JBTS is a relatively mild malformation and patients with other ciliopathies, such as Bardet-Biedl syndrome, can display learning difficulties and developmental delays without abnormalities on brain MRI. Together, this suggests that cilia may play additional, more subtle roles in CNS function beyond development. In this study, we investigated the shared and unique consequences of JBTS gene dysfunction on the zebrafish CNS. We describe multiple zebrafish mutants in JBTS genes which lack any gross morphological CNS abnormalities despite abnormal CNS primary and motile cilia. We further show a dysregulated expression of genes associated with neuronal function paralleled by abnormal behaviours. We therefore speculate a role for primary cilia in regulating the function of differentiated neurons and propose the models described here as powerful tools to study this further.

In this study, we identified that dysfunction of JBTS genes results in abnormal cilia throughout the brain. It is interesting to note that the loss of the different JBTS genes affects cilia differently in each mutant and with distinct local effects. In some cases, primary cilia were affected only in one brain region and in others motile cilia were more affected than primary cilia. These findings suggest tissue-specific roles for JBTS proteins in ciliary biology. The presence of cilia on differentiated neurons such as Purkinje or eurydendroid cells in the cerebellum supports their role in mature neurons, possibly in regulating the function of these neurons.

This is consistent with the transcriptomic analysis which identified an enrichment of gene sets associated with neuronal activity in all JBTS mutants. Genes encoding both excitatory and inhibitory neurotransmitter receptor subunits as well as voltage-gated potassium, sodium and calcium ion channel subunits were identified in these gene sets, all of which are fundamental for neurotransmission. As highly specialised signalling hubs, primary cilia are primed to transduce signalling pathways that regulate neurotransmission. For example, neuronal primary cilia are enriched in GPCRs that function in modulating neuronal activity. Primary cilia on cultured pyramidal neurons have been shown to modulate excitatory synapses via SSTR3, and cilia disruption leads to an accumulation of AMPA receptors (Tereshko et al. 2021). Primary cilia are also specialised Ca^2+^ signalling organelles. The ciliary Ca^2+^ channel PKD2L1 plays a role in regulating hippocampal excitability in mice, and mice lacking PKD2L1 were found to be more susceptible to epileptic seizures (Vien et al. 2023). Primary cilia on differentiated neurons within the zebrafish CNS could therefore concentrate receptors or channels that influence neuronal function. Future work will be required to analyse the signalling machineries present in primary cilia within the zebrafish CNS.

Interestingly, the transcriptomic analyses showed distinct trends between the different mutants, as we find some of the same genes to be downregulated in *cc2d2a^−/−^* but upregulated in *talpid3^−/−^*. This suggests distinct consequences of JBTS gene dysfunction on the expression of these neurotransmitter receptors and voltage-gated ion channels. As Cc2d2a and Talpid3 have different functions within the primary cilium, it is possible that their dysfunction has different consequences. For example, *cc2d2a* and *talpid3* dysfunction in the zebrafish causes retinal degeneration through two different mechanisms. In *cc2d2a^−/−^* larvae, retinal degeneration occurs due to disrupted vesicle fusion within photoreceptors while *talpid3*^−/−^ photoreceptors display defective basal body positioning and docking to the apical cell surface (Ojeda Naharros et al. 2018; Ojeda Naharros et al. 2017). Likewise, Cc2d2a and Talpid3 loss may also have distinct consequences within neurons, explaining the different behavioural phenotypes observed. The loss of posture we observe is also intriguing, as the cerebellum plays a very important role in postural control. Loss of posture is also observed in zebrafish lacking Arl13b, which have malformed cerebella (Zhu et al. 2020). However, it is challenging to directly link changes in larval behaviour to only changes in neuronal activity in the brain and/or spinal cord, given the additional body curvature and retinal degeneration phenotypes seen in *cc2d2a^−/−^*. We did however select zygotic *cc2d2a^−/−^* with mild curvature and with an inflated swim bladder and still observed clearly abnormal behaviour, suggesting that the curved tail is not entirely responsible for abnormal swimming. To better tease these two features apart, it would be interesting to analyse neuronal activity in JBTS mutants using genetically-encoded calcium indicators expressed within neurons. A recent preprint used this method to show that zebrafish entirely lacking primary cilia due to a mutation in the ciliary gene *ift54* show reduced photic-induced and spontaneous neuronal activity (D’Gama et al. 2023 preprint).

We observed similar transcriptional and morphological phenotypes in *bbs1* and JBTS mutants, despite the *bbs1* gene being causative for a distinct ciliopathy. JBTS is characterised by the MTS which is absent in BBS, but both JBTS and BBS patients can present with variable cognitive impairment that could be caused by neuronal circuit dysfunction. Therefore, though zebrafish JBTS mutants do not show a brain malformation comparable to the MTS, they may have neuronal circuit anomalies that are also present in zebrafish *bbs1* mutants, resulting in similar transcriptomic phenotypes.

The RNA sequencing analysis performed here carries some limitations which must be taken into account. The genetic heterogeneity of outbred zebrafish is a major confounder for differential gene expression analysis, and hierarchical clustering shows that the effect between paired samples (siblings) is greater than the effect between mutants and controls. We therefore used a paired statement and the siblings as controls to mitigate these effects. This study design may explain why we found so little overlap between the different mutants at the level of individual differentially expressed genes. Furthermore, we saw rather small effect sizes in our mutants, which is likely explained by the fact that we carried out RNA sequencing of whole larvae, which naturally dilutes tissue-specific effects. In light of this, we used enrichment analysis to find GO terms that were enriched at the extremes of the gene lists, ranked by P-value in ORA and log fold change in GSEA. The fact that we could find the same GO terms in all mutants validates the results of this enrichment analysis and highlights the CNS of zebrafish JBTS mutants as an interesting area for further investigation.

An important question raised by our data is why JBTS gene dysfunction in the zebrafish does not lead to structural CNS defects as in mice, especially given that we observe ciliary phenotypes in zebrafish JBTS mutants. One hypothesis could be that maternal contribution of mRNA and/or proteins in zygotic zebrafish JBTS gene mutants and retention of some neuronal cilia may obscure early developmental phenotypes. However, we performed our analyses on maternal zygotic mutants whenever possible and mz *cc2d2a^−/−^*, for example, have no morphological defects in the CNS while showing a strong reduction in Arl13b-positive cilia. We may therefore be seeing species-specific requirements for JBTS-associated genes in neurodevelopment. While many aspects of brain development are conserved between zebrafish and mammals, there are some notable exceptions, for example in cerebellar development. In mammals, Shh secreted by Purkinje cells is transduced by primary cilia present on granule cell precursors in the external granule layer (EGL), causing expansion of the granule cell precursor population and enabling formation of the mature granule cell layer (Spassky et al. 2008). However, the presence of an EGL in zebrafish has been debated. While an earlier study reported a lack of EGL in the developing zebrafish cerebellum, a more recent study could show the presence of proliferative cells in a region corresponding to the EGL (Chaplin et al. 2010; Biechl et al. 2016). However, in the zebrafish cerebellum, *shh* is not expressed by Purkinje cells but is instead expressed by *olig2+* eurydendroid cells, and whether eurydendroid cell-derived Shh is transduced by primary cilia on granule cell precursors during EGL development in zebrafish remains a mystery (Biechl et al. 2016). The lack of morphological defects in the CNS of zebrafish JBTS mutants may therefore be due to differences in the signalling pathways required during neurodevelopment or in the role or primary cilia in transducing these pathways. It is interesting that we observe no CNS morphological phenotypes in the analysed JBTS mutants while some zebrafish lacking Arl13b display a reduction in Purkinje cells and granule cells (Zhu et al. 2020). Given recent data suggesting that ARL13B may also function outside of the primary cilium in mice during cerebellar development, it could be that the cerebellar phenotypes observed in zebrafish *arl13b^−/−^* are due to a total loss of Arl13b in the cell compared to a reduction in Arl13b-containing cilia in the JBTS mutants (Suciu et al. 2021).

Signalling pathways may also be transduced differently by the primary cilium in mammals and zebrafish. For example, mouse mutants lacking cilia display ventral neural tube patterning defects similar to *Shh* mutants (Huangfu et al. 2003; Liu et al. 2005). Conversely, zebrafish mutants lacking cilia display dampened but expanded Hh signalling in the neural tube and somites due to Hh-independent Gli1 activity, at least at early stages (Huang and Schier 2009). Similarly, the primary cilium appears to be nonessential for Wnt signalling in the zebrafish while its role in mammals is disputed (Huang and Schier 2009). It is intriguing that our RNA sequencing analysis revealed no impairments in any signalling pathways, including Hh, in all JBTS mutants. However, this could be due to effects of whole tissue analysis. Shh is strongly expressed in the posterior mesenchyme of the fin bud and floor plate at 72 hpf in zebrafish (Fabien Avaron et al. 2013). These cells represent a small population compared to the whole larva, which could lead to a strong dilution effect. Additionally, we show that cilia are reduced but not completely lost in the brain of zebrafish JBTS mutants, potentially preserving some Hh signalling.

Taken together, the thorough and systematic analysis of multiple zebrafish JBTS gene mutants performed in this work has revealed a conserved role for the encoded proteins in building and/or maintaining cilia within the brain, but shows that their loss does not cause patterning, differentiation or proliferation defects. Rather, loss of JBTS proteins causes dysregulation of genes that are important for neural circuit function and results in abnormal behaviours. These models therefore offer the unique opportunity to study the importance of primary cilia-mediated signalling in neurons beyond early neurodevelopment.

## Methods

### Ethics statement

All animal protocols were in compliance with internationally recognised and with Swiss legal ethical guidelines for the use of fish in biomedical research. Zebrafish husbandry and experimental procedures were performed in accordance with Swiss animal protection regulations (Veterinäramt Zürich, Tierhaltungsnummer 150, Tierversuch ZH116/2021-33632).

### Animal husbandry

Zebrafish (Danio rerio) were maintained as described previously (Aleström et al. 2020). Zebrafish embryos were raised in embryo medium at 28°C and where necessary treated with phenylthiourea until 5 days post fertilisation (dpf) to prevent pigment development (Westerfield 2000). The *cc2d2a^uw38^*, *talpid3^i264^*, *cep290^fh297^, togaram1^zh510^*and *bbs1^ka742^* mutants were described previously (Ben et al. 2011; Owens et al. 2008; Latour et al. 2020; Masek et al. 2022; Lessieur et al. 2019). The *Tg(tagRFP-T:PC:GCaMP5G)* and *Tg(olig2:EGFP)* transgenic lines were described previously (Matsui et al. 2014; Shin et al. 2003). The *Tg(βactin:GFP-centrin)* line was a kind gift from Brian Ciruna. The *Tg(ubi:Arl13b-mCherry)* zebrafish line was established by a conventional Tol2-mediated transposition method (Suster et al. 2009). The *arl13b* coding sequence from the Drummond group’s ubiquitin:arl13b-GFP construct (Austin-Tse et al. 2013) was inserted into a pME vector and subsequently recombined with a p5E vector containing the ubiquitin promoter and a p3E-mCherrypA construct, all integrated into a pDESTTol2CG2 vector. Subsequently, the construct was injected into embryos at the one-cell stage, and a founder fish was identified through the expression of the green heart marker indicative of the transgene. Outcrosses were performed for more than 3 generations, and Mendelian ratios of transgene positive offspring indicate a single insertion site for the transgene. Lines are available upon request by contacting the corresponding author.

### Generation of *inpp5e* mutant

sgRNAs for CRISPR/Cas9 mutagenesis were designed with chopchop (https://chopchop.cbu.uib.no/): sgNRA-*inpp5e*-ex1-1 (5‘-GGAGAAGAGGGCGGTCGGAG-3‘) and sgRNA-*inpp5e*-ex1-2 3’-GGGGTTCAGAACGCTATATG-5’ targeting exon1 of *inpp5e*. sgRNAs were mixed with Cas9 protein (M0646M NEB, or B25641 invitrogen) and coinjected into one-cell stage embryos using a microinjector (Eppendorf). Amplification of the target regions for genotyping was performed using primer pairs 5‘-AGGCACGTATCCTCTTCTGG-3‘ and 3‘-CAAGACGACACATCAGCACA-5‘ for exon 1 in *inpp5e*. The *inpp5e^zh506^* line was established from an F1 fish harbouring two deletions in exon 1: one 29 bp deletion at the target site of sgNRA-*inpp5e*-ex1-1 and a second 9 bp deletion at the target site of sgNRA-*inpp5e*-ex1-2; together this results in a 38 bp deletion visible on the electrophoresis gel. The first 29 bp deletion leads to a premature termination codon after 1 aa (NM_001102619:c.648del29,739del9, p.S217del9Fs*1) (Fig. S1). F1 fish were outcrossed for at least 3 generations.

### RNA sequencing

#### Sample Collection

Mutant zebrafish females were crossed with heterozygous males to generate maternal zygotic mutants and their heterozygous sibling controls. The fish embryos were staged and raised in E3 until 72 ± 1 hours post-fertilisation. Following this, the larvae were euthanized in ice-cold 0.4% MS-222 solution, and a small tail biopsy was obtained to genotype the larvae, while the remaining body was preserved in RNAlater. Siblings sharing the same genotype were pooled, with a minimum of 10 bodies per pool. Total RNA was extracted from these pooled samples using the RNeasy Plus Micro Kit (Qiagen, USA). For each zebrafish mutant line, three independent pairs of samples were collected, each consisting of pooled mutants (−/−) and their sibling controls (+/−) from the same clutch. In the case of *talpid3*, where adult mutants cannot be raised, a heterozygous incross was performed to obtain zygotic mutants (−/−) and sibling controls (+/+).

#### Library Preparation and Sequencing

Library preparation and sequencing was performed at the Functional Genomics Center Zurich (FGCZ) of University of Zurich and ETH Zurich. The quality of the isolated RNA was determined with a Fragment Analyzer (Agilent, Santa Clara, California, USA). Only those samples with a 260 nm/280 nm ratio between 1.8–2.1 and a 28S/18S ratio within 1.5–2 were further processed. The TruSeq Stranded mRNA (Illumina, Inc, California, USA) was used in the succeeding steps. Briefly, total RNA samples (100-1000 ng) were poly A enriched and then reverse-transcribed into double-stranded cDNA. The cDNA samples were fragmented, end-repaired and adenylated before ligation of TruSeq adapters containing unique dual indices (UDI) for multiplexing. Fragments containing TruSeq adapters on both ends were selectively enriched with PCR. The quality and quantity of the enriched libraries were validated using the Fragment Analyzer (Agilent, Santa Clara, California, USA). The product is a smear with an average fragment size of approximately 260 bp. The libraries were normalized to 10nM in Tris-Cl 10 mM, pH8.5 with 0.1% Tween 20. The Novaseq 6000 (Illumina, Inc, California, USA) was used for cluster generation and sequencing according to standard protocol. Sequencing were paired end at 2 X150 bp or single end 100 bp. Raw sequencing data are deposited on the NCBI Sequence Read Archive (SRA). The data can be found with the BioProject ID: PRJNA1070872” (http://www.ncbi.nlm.nih.gov/bioproject/1070872)

#### Data Analysis

The generated reads were mapped to the zebrafish reference sequence GRCz11 using STAR aligner (version 2.7.10) and ENSEMBL v103 annotations. Gene feature counts were obtained, and descriptive statistics were employed for sample comparisons. Sample-wise Pearson correlation analysis was conducted on the rlog-transformed counts, along with principal component analysis using the top 25% most variable z-transformed genes via PCAtools (version 1.12.0) in R.

Differential expression analysis was carried out on a line-by-line basis using DESeq2 (version 1.40.1), adding the paired feature to the regression model. Genes with an adjusted P-value < 0.05 were considered significantly differentially expressed. Enrichment analysis was performed using the R package clusterProfiler (version 4.9.0) using genes with a threshold of P-value < 0.01 and an absolute fold change > 1.5 for the overrepresentation analysis. Ontology terms with an adjusted P-value < 0.05 were considered significantly enriched in the gene set enrichment and overrepresentation analysis. To account for highly similar terms, GO semantic similarity analysis was performed using GOSemSim (version 2.26.0). The results were visualized using the R packages ggplot2 (version 3.4.2), UpSetR (version 1.4.0), and Pheatmap (version 1.0.12).

### Immunohistochemistry

Whole larvae were fixed in 4% paraformaldehyde (PFA), 80% MeOH in DMSO or 2% trichloroacetic acid (TCA) at room temperature or 4°C (for details, see Table S9) and washed in PBS. For larvae fixed with 4% PFA, permeabilisation was carried out using acetone at −20°C. Larvae were blocked using PBDT (PBS + 1% BSA + 0.5% Triton X-100 + 1% DMSO) with 10% goat serum for 30 minutes at room temperature. Larvae were incubated with primary antibodies overnight at 4°C, washed in PBDT and incubated in secondary antibodies for at least 2 hours at room temperature or overnight at 4°C (see Table S9). Nuclei were counterstained for 15 minutes using 1μg/mL DAPI in H2O. Refractive index matching was carried out in 70% glycerol in PBS and larvae were mounted for imaging using Mowiol containing DABCO. Confocal images were acquired using the Leica TCS LSI, Leica STELLARIS 5 or Olympus Spinning Disk confocal microscopes. Widefield images were acquired as z-stacks using the Leica THUNDER Imager Model Organism fluorescence microscope before processing using parallax correction followed by large volume computational clearing to remove background with Leica LAS X software.

### CLARITY whole brain clearing and light-sheet imaging

The CLARITY tissue clearing method was carried out according to the previously published protocol (Chung and Deisseroth 2013). Briefly, adult fish were euthanised in ice cold water followed by decapitation. Brains were dissected and fixed in 4% PFA for 3 hours at room temperature. Brains were incubated in hydrogel monomer solution (4% acrylamide + 0.25% VA-044 in PBS) for 12-24 hours at 4°C, rocking and then incubated for 3 hours at 37°C, without rocking. Passive clearing was carried out using CLARITY clearing solution (4% SDS in 200mM boric acid) for 2-3 weeks at RT with gentle rocking, with fresh CLARITY clearing solution exchanged every 1-3 days. After clearing, brains were washed with 0.1% PBST (PBS + 0.1% Triton X-100). Nuclei were counterstained with 1μg/mL DAPI in H2O overnight at RT. Brains were embedded in 1.5% low-melting agarose for imaging and refractive index matching was carried out using ∼88% Histodenz. Samples were imaged using the mesoSPIM light-sheet microscope (mesoSPIM V6 “Revolver”) (Voigt et al. 2019; Vladimirov et al. 2023 preprint). For mesoSPIM imaging, the embedded samples were mounted inside a quartz cuvette (Portmann Instruments UQ203, 45 x 12.5 x 7.5 mm, filled with ∼88% Histodenz). The sample cuvette was dipped into an immersion chamber (Portmann Instruments UQ-753, 40 x 40 x 50 mm) filled with index-matching oil (1.46, Cargille 50350). The samples were imaged with a 5X objective and a z step size of 2 μm.

### Behavioural Analysis

Zebrafish locomotion was analysed using an automated Zebrabox infrared tracking system (ViewPoint Life Science, Lyon, France). 3 dpf or 6 dpf larvae were placed in a 96-well plate containing embryo medium. Locomotion was recorded for three consecutive periods of 5 minutes without light, 5 minutes with light and 5 minutes without light and the distance that a single larva moved was recorded during 10 second intervals.

The zebrafish touch-evoked escape response was measured at 2 dpf using the Leica THUNDER Imager Model Organism fluorescence microscope with a frame rate of 0.05 seconds. Larvae were manually dechorionated at least 1 hour before recording. Larvae were transferred to a petri dish containing embryo medium one minute before initiating the response and allowed to habituate. The touch-evoked escape response was initiated by lightly touching the tail of the larva with a metal poker. Displacement was quantified as the distance between the head position in the first frame preceding the touch-evoked escape response and the head position in the third frame after the touch-evoked escape response.

Graph plotting and statistics were carried out using GraphPad Prism. For quantifications, data were pooled from at least 1-2 independent experiments (independent clutches/independent experiment days) containing multiple animals. Larval zebrafish sex is not determined at this stage.

### Imaging Data analysis

Quantification of Purkinje and eurydendroid cell number was carried out using the data analysis platform Dataspace, software available at https://github.com/skollmor/dspace (Kollmorgen et al. 2020). Image stacks encompassing the Purkinje and eurydendroid cells in one cerebellar hemisphere were pre-processed by applying image convolution and thresholding to remove background. Cells within each image slice were then identified based on the presence of fluorescent local maxima (“detections”) and overlapping detections in each slice consolidated to a single identified cell (Fig. S4).

The Vglut1-positive fluorescence area was quantified using Fiji. Sum intensity z projections were generated from image stacks encompassing the Vglut1+ presynaptic terminals in one cerebellar hemisphere. These sum intensity z projections were converted to 16-bit before automatic thresholding using the Huang method. The area of the thresholded region was then calculated. This area was normalised to the size of the larva’s head, calculated by multiplying the length of the head from the anterior edge of the eyes to the most posterior point of the head with the width of the head just posteriorly to the eyes.

The width of eurydendroid cell tracts was quantified using Fiji. Maximum intensity z projections were generated from image stacks encompassing the eurydendroid cell commissure. The width was measured at a position at the left, centre and right of the tract and the three widths averaged to calculate the eurydendroid cell tract width.

The area of the forebrain, midbrain and hindbrain was quantified using Fiji. Polygonal ROIs were manually drawn around the whole brain, forebrain, midbrain and hindbrain on single slice images of the brain. The area of the forebrain, midbrain and hindbrain ROIs were measured and normalised to the area of the whole brain ROI.

Images were prepared using Fiji image processing software and videos were prepared using Imaris image analysis software. For primary cilia images in Fig. 3C-H, eurydendroid cell axonal tract images in Fig. 4M-O and high resolution axon and synapse images in Fig. 6J-L, deconvolution was applied using Huygens Deconvolution software. For all quantifications, data were pooled from at least 2-3 independent experiments (independent clutches/independent experiment days) containing multiple animals (approximately n = 10 larvae per condition, minimum n = 5 larvae per condition). All measurements were made blind to the genotype. All means represent mean±s.d (standard deviation). Graph plotting and statistics were carried out using GraphPad Prism. Experiments on juvenile fish included both sexes, while in larval zebrafish sex was not determined.

## Acknowledgements

We thank the Functional Genomic Center Zurich (FGCZ) for conducting the RNAseq experiment as well as the Center for Microscopy and Image Analysis Zurich (ZMB) for their support. Special thanks go to Brian Ciruna, Iain Drummond, Masahiko Hibi and Reinhard Köster for sharing fish lines and/or reagents. ARN was supported by the University Research Priority Program (URPP) ‘Adaptive Brain Circuits in Development and Learning (AdaBD)’ of the University of Zurich. This work was supported by SNSF grants P00P3_170681 and PP00P3_198895/1 to RBG.

